# The nature and intensity of mechanical stimulation drive different dynamics of MRTF-A nuclear redistribution after actin remodeling in myoblasts

**DOI:** 10.1101/395962

**Authors:** Lorraine Montel, Athanassia Sotiropoulos, Sylvie Hénon

## Abstract

Serum response factor and its cofactor myocardin-related transcription factor (MRTF) are key elements of muscle-mass adaptation to workload. The transcription of target genes is activated when MRTF is present in the nucleus. The localization of MRTF is controlled by its binding to G-actin. Thus, the pathway can be mechanically activated through the mechanosensitivity of the actin cytoskeleton. The pathway has been widely investigated from a biochemical point of view, but its mechanical activation and the timescales involved are poorly understood. Here, we applied local and global mechanical cues to myoblasts through two custom-built set-ups, magnetic tweezers and stretchable substrates. Both induced nuclear accumulation of MRTF-A. However, the dynamics of the response varied with the nature and level of mechanical stimulation and correlated with the polymerization of different actin sub-structures. Local repeated force induced local actin polymerization and nuclear accumulation of MRTF-A by 30 minutes, whereas a global static strain induced both rapid (minutes) transient nuclear accumulation, associated with the polymerization of an actin cap above the nucleus, and long-term (hours) accumulation, with a global increase in polymerized actin. Conversely, high strain induced actin depolymerization at intermediate times, associated with cytoplasmic MRTF accumulation.

---

The development and differentiation of cells are known to be influenced by mechanical cues, such as deformation, stress, or the rigidity of the substrate [reviewed in ^1^]. In particular, changes in the mechanical properties of the surrounding matrix or tissue can alter cellular fate in diseases such as fibrosis, atherosclerosis, and cancer [^2^]. During development, the interplay between forces and deformation of the tissue organizes the embryo through gastrulation or neural closure [^3^], highlighting the importance of understanding the mechanics of morphogenesis.

At the cellular level, the influence of mechanical cues have been studied on a broad scale, ranging from that of single proteins, where cryptic sites can be unraveled by traction forces [reviewed in ^4^], to the global cellular scale, with the reorganization of the lamellipodium during motion [reviewed in ^5^] or the realignment of stress fibers in response to cyclic strain [^6^,^7^].

The main component of the mechanical signal converges towards the actin cytoskeleton, as it is the central agent of mechanical organization. Indeed, the mechanical properties of cells, such as their rheology and mechanosensitivity, primarily rely on their cytoskeleton, which is highly dynamic as the result of the constant interplay between cytoskeletal proteins and the mechanical environment. Adhesion proteins, which cross-talk with the actin cytoskeleton, are at the forefront of the mechanosensory apparatus of the cell. At the other end of the mechano-transduction pathways, the activity of transcription factors, such as YAP/TAZ and MRTF/SRF, have been shown to be mechanically dependent [reviewed in ^8^].

Serum response factor (SRF) [^9^] and its cofactor myocardin-related transcription factor (MRTF) [^10^, ^11^] are actin-sensitive transcription factors that regulate diverse biological functions, such as neural development [^12^,^13^], the circadian clock [^14^], fibroblast to myofibroblast transition [^15^,^16^], and muscle differentiation [^17^]. Together, they control the expression of hundreds of genes, especially those of the cytoskeleton, such as the actins [^18^]. The MRTF/SRF pathway has been shown to be a crucial regulator of muscle homeostasis in response to mechanical cues: it is required to increase muscle mass in response to mechanical overload, and its decreased activity during the lack of mechanical activity (or disuse atrophy) participates in muscle wasting [^19^,^20^].

The regulation of SRF by MRTF is controlled by the localization of MRTF in the cell: SRF is located in the nucleus where it can bind DNA, whereas MRTF is able to shuttle between the cytoplasm and the nucleus [^21^,^22^]. However, the bipartite nuclear localization signal (NLS) of MRTF is embedded in three G-actin-binding RPEL motifs [^23^,^24^,^25^]. Thus, the NLS is accessible only when actin is not already bound. Indeed, the availability of actin monomers determines accessibility of the NLS and the intracellular localization of MRTF; when monomers are abundant, the NLS is hidden and MRTF is sequestrated in the cytoplasm, whereas when they are scarce, MRTF can be imported into the nucleus and bind SRF [^22^,^26^]. Nuclear actin monomers can also prevent SRF binding, even when MRTF-A accumulates in the nucleus. Thus, the localization of actin and its polymerization state in the nucleus are also of importance: over-expression of the actin-NLS [^22^] blocks MRTF-A from binding SRF, whereas serum-stimulation induces nuclear actin polymerization through RhoA and mDia [^27^] and MRTF-A nuclear accumulation, as does nuclear actin expulsion by MICAL-2 [^28^].

The dependence of SRF/MRTF activity on actin dynamics was originally studied in response to biochemical cues, such as serum [^9^,^21^,^22^] and growth factor stimulation [^15^,^29^], which activate actin polymerization through the RhoA pathway and the two effectors ROCK [^30^,^31^] and mDia [^32^,^27^]. However, any signaling pathway that changes the balance between monomeric and filamentous actin can potentially alter MRTF localization. In particular, actin polymerization pathways regulated by small GTPases have been shown to be activated in response to mechanical cues [^30^,^33^]. Changes in the cellular geometric constraints through micropatterning [^15^] or changes in the rigidity of the substrate [^34^] can alter the F/G actin ratio in fibroblasts and epidermal cells and activate the nuclear localization of MRTF-A and the transcription of SRF target genes. The application of a constant global strain for 24 h also induces the transcription of the SRF targets in vascular smooth muscle cells [^31^], as does application of a cyclic strain to cardiomyocytes [^33^]. In fibroblasts, a constant force applied through collagen-coated microbeads activates the two RhoA downstream effectors ROCK [^30^] and mDia [^35^], and the target genes are activated within hours. More recently, Iyer *et al.* [^36^] investigated a very rapid dynamic regime (shorter than 5 min) in live HeLa cells and observed rapid actin polymerization and MRTF-A accumulation upon global force stimulation. All but the last of these studies where performed on fixed samples, with only the last two [^30^, ^36^] exploring the dynamics of the system and finding very different response times (from 5 to 50 min) for MRTF-A nuclear accumulation and actin polymerization.

Here, we assessed the dynamic responses of actin and MRTF-A to two different types of mechanical stimulation, local or global, over a large range of time scales, from a few minutes to two hours, and investigated the unexplored question of the role of strain level. We used myoblasts, a cell type that has seldom been studied in the context of MRTF-A/SRF, whereas this pathway is central to the adaptation of skeletal muscle to force. We observed a strong correlation between actin polymerization and relocation of MRTF-A into the nucleus. First, we showed that the location of MRTF-A-GFP within myoblasts correlates with its expression level and the levels of G- and F-actin. Second, we used custom-built magnetic tweezers to show that a force applied through a single bead can trigger the local polymerization of actin around the bead and relocation of MRTF-A to the nucleus within 30 min. Finally, we used a stretching device that applied controlled strains and observed nuclear relocation of MRTF-A under moderate strain at two timescales: a few minutes, and a few tens of minutes. We linked those two regimes to the reorganization of two different parts of the actin cytoskeleton, apical and basal stress fibers. Under higher strain, we showed that the first rapid response is maintained, though the actin cytoskeleton is later disrupted and MRTF-A relocates to the cytoplasm.

## Materials and methods

### Cell Culture

C2C12 murine myoblasts from the ATCC were grown in DMEM with 10% FCS, 1% penicillin, and streptomycin at 37°C and 5% CO2. DMEM with red phenol was replaced by DMEM without red phenol the day before the experiments to allow optimal fluorescence imaging.

### Transfection and live markers

SiR-actin (Spirochrome) was added to cells at a concentration of 50 nM approximately 15 h before the experiment and used without rinsing. DAPI was added to live cells 30 min before the experiment. The MRTF-A-GFP plasmid has been described previously [^22^]. The mCherry-actin and LifeAct-mCherry plasmids were kind gifts of Maïté Coppey (Institut Jacques Monod, Paris, France) and Claire Hivroz (Institut Curie, Paris, France).

For all experiments, except those with SiR-actin, cells were transfected using Nanofectin (PAA Laboratories, Pasching, Austria). Approximately 110,000 cells were transfected 18 h before the experiment, following the manufacturer’s instructions, with 0.75 to 2.5 µg DNA and incubated for 6 h with 0.75 to 2.5 µg Nanofectin. Co-transfection with mCherry-actin was performed by adding 1 µg mCherry-actin DNA and 1 µg Nanofectin and following the usual transfection protocol. After the end of commercialization of Nanofectin, transfections were performed using Lipofectamine 3000 (Invitrogen), using the same amount of DNA and following the manufacturer’s instructions. Lipofectamine and DNA were incubated with the cells for 24 h and left in the culture medium throughout the experiment.

### Immunostaining

Cells were fixed in a 4% paraformaldehyde solution for 20 min and stained with phalloidin Alexa Fluor 647 or 488 at 0.026 nmol/l and DAPI at 1 μg/ml (all three from Life Technologies) for 30 min at room temperature or overnight at 4°C, after permeabilization with 0.5% Triton X-100 in PBS and saturation. Staining of MRTF-A in un-transfected cells was performed with anti-MRTF-A antibody H140 (Santa Cruz Biotechnologies), at 0.8 μg/ml for 30 min at RT, and anti-rabbit Alexa Fluor 488 (Life Technologies) at 4 µg/ml for 30 min at RT.

### Magnetic Tweezers

The custom built magnetic tweezers are based on an electromagnet (66.5 mm long coil of approximately 800 turns of 0.5 mm copper wire, forming 8 layers) placed around a cylindrical soft-iron core (5.10 mm in diameter, 144 mm long) with a 60° cone-shaped tip (see SI Fig. 1). The electromagnet provided up to 1.2 A through a home-made current generator controlled by a function generator (TG1010, TT Instruments). The magnetic tweezers were used to apply local forces to cells through adhesive 4.5-µm super-paramagnetic beads (Dynabeads M450 Epoxy Invitrogen). The force applied to a bead depends on the current provided to the coil and the distance from the bead to the tip. The forces were pre-calibrated by suspending the Dynabeads in liquid polydimethylsiloxane (PDMS) of known viscosity; for each current provided to the electromagnet, the bead velocity *versus* the distance to the tip was measured by analyzing the recorded trajectories using Stokes law. A force *versus* distance calibration was obtained for each current.

**Figure 1.**
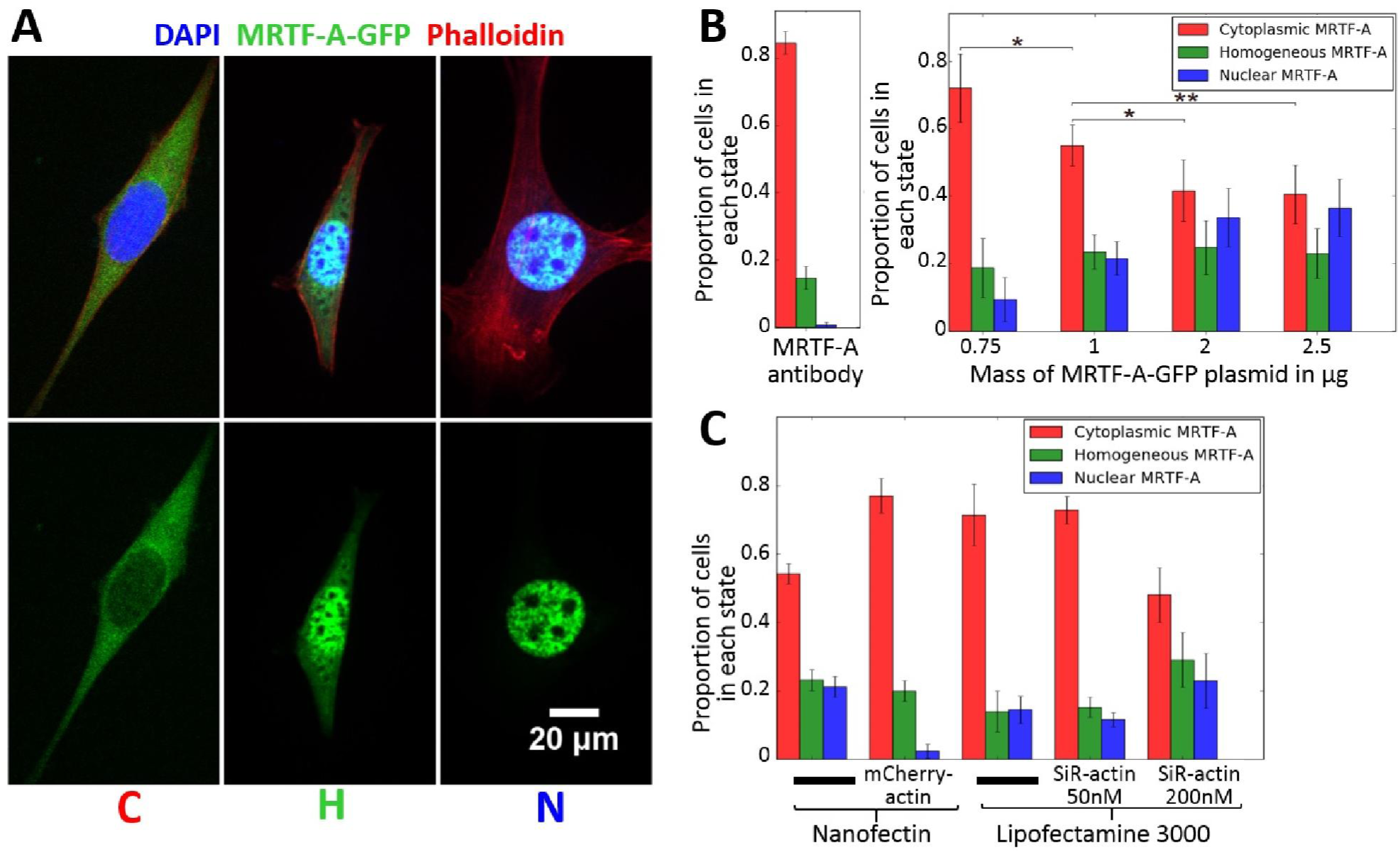
A.Examples of cells classified according to the intracellular localization of MRTF-A GFP, from left to right: mainly nuclear MRTF-A, mainly cytoplasmic MRTF-A, and homogeneously distributed. B.Localization of MRTF-A GFP as a function of the mass of MRTF-A GFP plasmid DNA used for transfection with Nanofectine. More DNA leads to nuclear accumulation of MRTF-A due to its overexpression relative to that of the G-actin pool (0.75 µg: 75 cells, 1.0 µg: 260 cells, 2.0 µg: 113 cells, 2.5 µg: 126 cells). The reference state for no transfection was obtained using anti-MRTF-A antibodies C.Localization of MRTF-A GFP as a function of transfection reagent, actin overexpression, and the F-actin stabilizing fluorescent marker SiR-actin.

For magnetic tweezer experiments, the beads were coated with fibronectin (5µg fibronectin for 4.10^7^ beads for 30 min at 37°C), then saturated with 10 µg/mL BSA for 30 min at 37°C. Cells were seeded on 22 × 22 mm glass coverslips coated with fibronectin (5 µg/mL in DMEM for 30 min at 37°C), 24 h before the experiment. Thirty minutes before the experiment, a suspension of fibronectin-coated beads was added to the cells and left to incubate for 30 min. Just before an experiment, the non-attached beads were removed by gentle rinsing, to avoid accidental mechanical stimulation at this step, and then the coverslip was mounted under the microscope for observation (DM IRB, Leica, Wetzlar, Germany, 100X oil immersion objective, 1.25 NA or 40X air objective, 0.70 NA). The electromagnet and core were mounted on a micro-manipulator (Inject-Man NI2, Eppendorf) at a 45° angle to the microscope stage (SI Fig. 2). The axis of the core was aligned with the center of the observation zone. All reported experiments were performed at a distance of 280 µm from the bead to the tip. At this distance, the maximum force that could be applied to a single bead, with the maximum 1.2 A current in the electromagnet, was approximately 1 nN.

**Figure 2.**
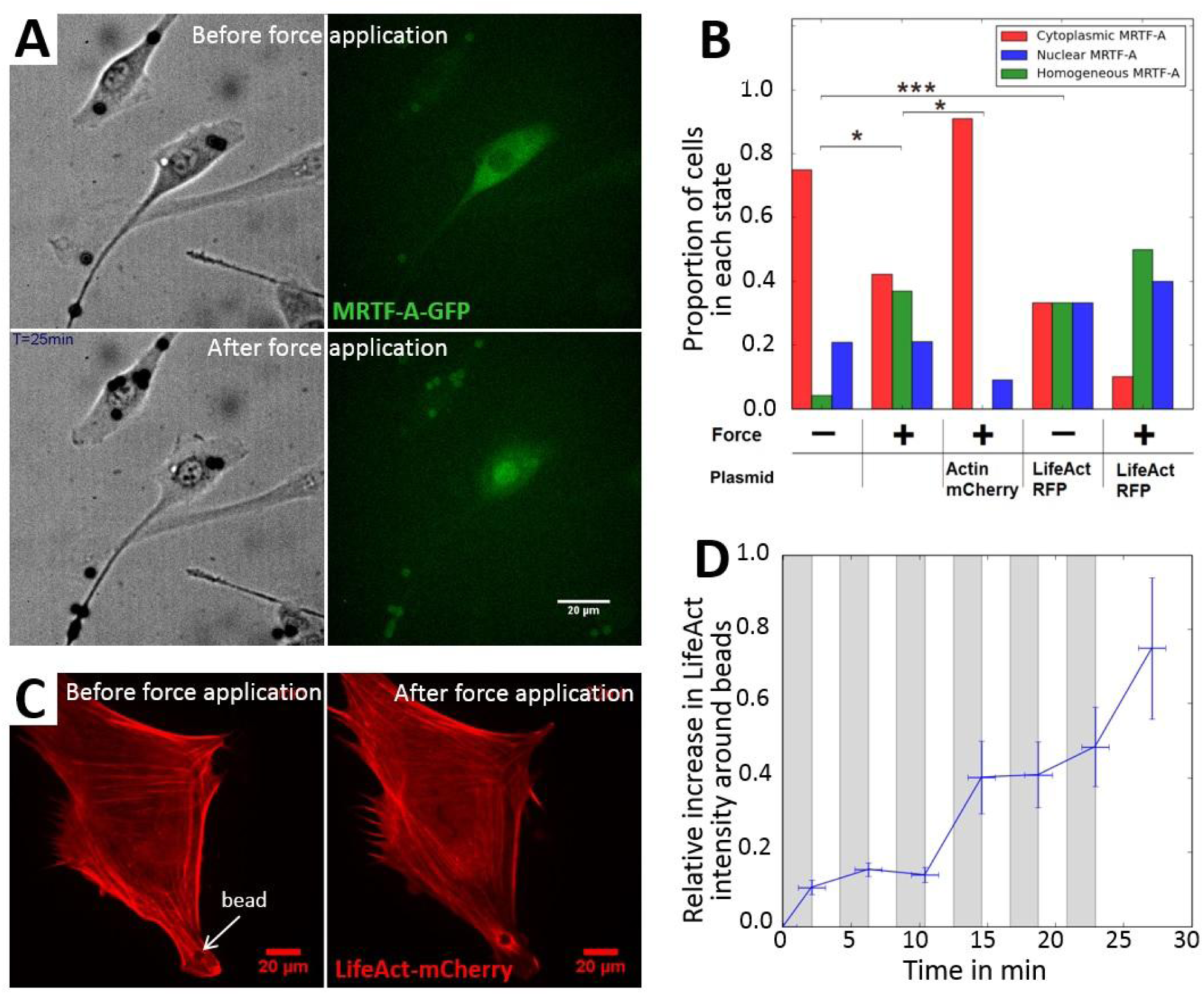
Fibronectin-coated beads actuated by magnetic tweezers induce local actin reorganization and global re-localization of MRTF-A to the nucleus. A.Live images of a cell with attached beads before the application of force, and after six applications. By the end of the experiment, MRTF-A-GFP relocated to the nucleus. B.Final state of MRTF-A-GFP localization in cells subjected to a 1-nN force through adherent fibronectin-coated microbeads. The force was applied six times for 125 seconds. Number of cells: 29 for the control, 23 with force, 11 with mCherry-actin and force, 10 for control with LifeAct- mCherry, and six for force with LifeAct-mCherry. *p* values were calculated using Fisher’s exact test (\**p* < 0.05, \*\**p* < 0.01, \*\*\**p* < 0.0001). C.Images of the actin cytoskeleton with LifeAct-mCherry before and at the end of the application of force. An actin ring is clearly visible around the bead at the end of the experiment. D.Relative increase in the mean fluorescence intensity of LifeAct-mCherry per pixel in the vicinity of a bead (see SI Figure 5) relative to the fluorescence mean intensity per pixel for the whole cell when subjected to repeated force (six cells). The zones in grey correspond to the application of force and the zones in white to the release of force.

### Cell Stretcher

Stretching experiments were performed using a custom-built device (SI Fig. 2) that allowed the cells to be visualized under the microscope while stretching them. Twenty-four hours before an experiment 110,000 cells were seeded on a PDMS disk (30 mm in diameter, 0.3 mm thick, PDMS + 10% curing agent from Sylgard Silicon Elastomer) coated with fibronectin (5 µg/mL in DMEM for 30 min at 37°C). The PDMS disk was mounted between two cylinders. The assembly was placed, with the side on which the cells were seeded face down, in a cylindrical tank which contained culture medium supplemented with 1.5% HEPES. The bottom of the vessel consisted of a glass coverslip 30 mm in diameter to allow observation of the cells under an inverted microscope. The PDMS disk was stretched by pushing a cylindrical transparent plastic post and thus the cells seeded on it were also stretched. The distance between the initial position of the PDMS disk and the final position after pushing the post determined the strain imposed on the disk, which was equal to the relative increase in the surface of the stretched area. Calibration using a PDMS disk micro-patterned with fluorescent fibronectin confirmed a uniform radial strain. The measured deformation was also in good agreement with the calculated deformation (see SI).

For live experiments, the experimental chamber was mounted on the motorized stage (Prior ProScan II) of an inverted microscope (Olympus IX81) and enclosed in a thermalized box (The Cube2, Life Imaging). The desired strain was then applied in less than 5 s at the initial time and kept constant over time. During the first 20 min of the experiment, the sample was scanned to locate cells expressing MRTF-A-GFP and their position was marked. Every 10 to 15 min, a new image of each recorded position was taken, making it possible to follow each cell over time. At the end of a live experiment, the sample could be fixed for later labeling and imaging of the final state.

### Classifying the cells according to MRTF-A-GFP localization

Cells expressing MRTF-A-GFP were classified according to the major localization of MRTF-A-GFP in the cell, as illustrated in Fig 1A. Cells for which the nucleus was clearly visible and bright in the MRTF-A-GFP channel were labeled as those with mainly nuclear MRTF-A (“Nuclear MRTF-A”, in blue in the graphs). On the contrary, cells for which the nucleus was clearly visible and dark in the green channel were labeled as those with mainly cytoplasmic MRTF-A (“Cytoplasmic MRTF-A”, in red in the graphs). Cells for which the border between the nucleus and cytoplasm could not be distinguished in the green channel were labeled as “Homogeneous MRTF-A” (in green in the graphs).

### Quantitative fluorescence analysis

Area, mean intensity, and integrated density of the fluorescence were measured on images with the ImageJ processing program, for three regions of each cell: the whole cell, the nuclear region, and a perinuclear region of approximately the same thickness as the nuclear region. The perinuclear region was drawn by hand on the images, the nuclear region determined by thresholding on the DAPI channel, and the whole cell region determined by thresholding on the GFP channel of individual cells and drawn by hand when separating adjacent cells was necessary.

### Database organization and sub-group selection

Data concerning the experiments were stored in an SQL database to allow data storage for a large number of cells in an easily accessible framework. The parameters of each experiment, such as the date, strain rate, or starting time of observation, were stored in the Experiment table. Parameters depending on the field of view, such as the total number of cells on the image, were stored in the Zone table. Parameters measured on individual cells, such as plasmid expression, main localization of MRTF-A, or times when MRTF-A localization changed, were stored in the Cell table. Each cell from this table belonged to a field of view identified in the Zone table, and each field of view belonged to an experiment from the Experiment table. This data structure allowed us to select groups of cells following a large variety of queries: for example, all cells expressing actin-mCherry with no more than 25 cells in the field of view, stretched at 10% strain rate, for which the initial state was “mainly cytoplasmic MRTF-A” and changed state between 10 and 30 min after the start of stretching.

### Statistical Analysis

Differences between the numbers of cells in the three categories of MRTF-A-GFP localization were statistically tested, either using a G-test of independence (similar to a chi-square test of independence) or Fisher’s exact test of independence, when the number of cells in one of the categories was too small. A difference was considered significant when the *p*-value was < 0.05 and values were corrected for multiple comparisons when necessary.

## Results

### The subcellular localization of MRTF-A correlates with its expression and the levels of G- and F-actin

We assessed MRTF-A localization during live experiments using C2C12 myoblasts transfected with a plasmid encoding MRTF-A-GFP [described in ^22^]. The subcellular localization of MRTF-A-GFP was observed by fluorescence microscopy and the cells were classified into three categories depending on the localization of MRTF-A-GFP (see Materials and Methods): mainly nuclear (“N”), mainly cytoplasmic (“C”), or homogeneously distributed (“H”), as illustrated in Figure 1A.

The localization of endogenous MRTF-A in non-transfected cells was assessed through immunostaining: it was mainly cytoplasmic for more than 85% of the cells, as expected from the literature [^21^] (Figure 1B and SI Figure 3).

**Figure 3.**
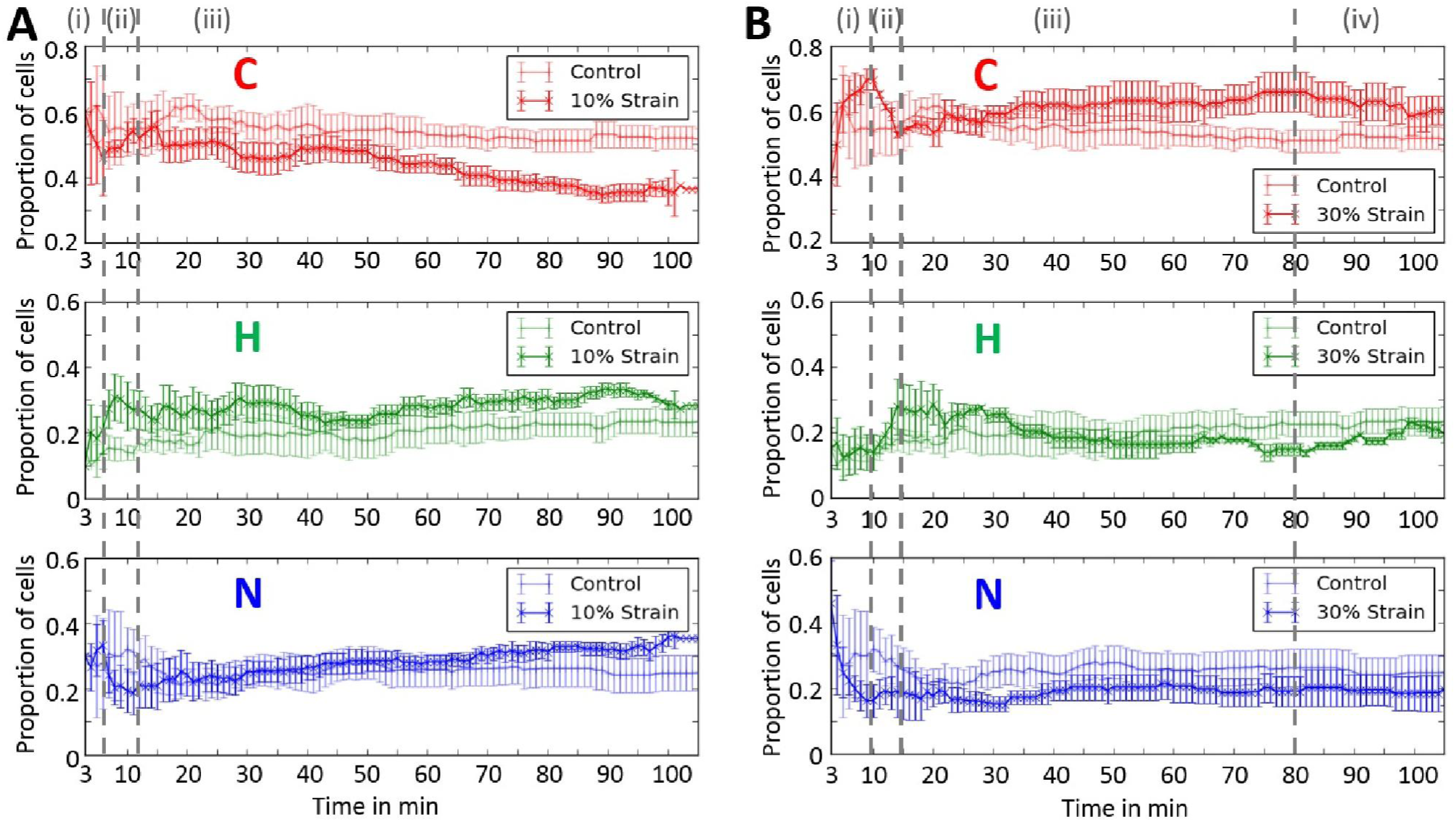
Repartition of the cells among the three categories of MRTF-A-GFP localization: “C” for mainly cytoplasmic, “N” for mainly nuclear, and “H” for homogeneously distributed. The cells were followed under a microscope over time, t = 0 corresponds to the beginning of stretching. A Cells stretched by 10% *vs* control (control: 143 cells, n = 5 independent experiments; 10% strain: 145 cells, n = 5 independent experiments). B Cells stretched by 30% *vs* control (control: 143 cells, n = 5 independent experiments, 30% strain: 101 cells, n = 3 independent experiments).

Various amounts of plasmid were used for transfection, ranging from 0.75 to 2.5µg of DNA for 110,000 cells. There was always less cytoplasmic MRTF-A-GFP in transfected than non-transfected cells, ranging from 70% of “cytoplasmic” cells for the lowest concentration of plasmid, down to 40% for the highest (Figure 1B): thus, the more plasmid used for transfection, the more MRTF-A accumulates in the nucleus. This is consistent with known mechanisms for the regulation of intracellular localization of MRTF-A through actin-binding. When MRTF-A is overexpressed due to multiple plasmid copies, there is insufficient G-actin to bind it and maintain it in the cytoplasm. The excess MRTF-A is thus free to accumulate in the nucleus. Even the lowest concentration of plasmid tested here caused an increase in nuclear localization of MRTF-A. For all subsequent experiments, 1 µg of plasmid was used, as a compromise between sufficient transfection efficiency and low-level overexpression.

Transfections were also performed using Lipofectamin 3000 with the same dose of plasmid per cell (1 µg for 110,000 cells) as for Nanofectin. Under the same experimental conditions, the localization of MRTF-A-GFP was more cytoplasmic than when using Nanofectin (Figure 1C), while the number of MRTF-A-GFP positive cells was at least doubled from 11% to more than 23%. Consistent with the dose-response experiment, the difference can be explained by the fact that Lipofectamin 3000 delivers the same amount of DNA to more cells than Nanofectin, each receiving fewer copies, resulting in lower overexpression and less nuclear accumulation of MRTF-A-GFP.

Co-transfection with mCherry-actin and MRTF-A-GFP expression vectors increased the levels of available G-actin to bind MRTF-A. Total actin levels (both G- and F-actin) increased under these conditions. The localization of MRTF-A-GFP in cells expressing both plasmids was very similar to that in non-transfected cells (Figure 1C). Thus, MRTF-A-GFP is maintained in the cytoplasm when the total amount of actin in the cell increases.

The G-actin / F-actin ratio was altered using SiR-actin (see Materials and Methods), a fluorescent probe for F-actin derived from the actin-stabilizing drug jasplakinolide. SiR-actin caused an increase in the nuclear localization of MRTF-A-GFP when used at a concentration of 200 nM (Figure 1C). The pool of G-actin was probably depleted by the stabilizing effect of SiR-actin on the filaments, thus favoring the accumulation of MRTF-A in the nucleus. SiR-actin had no effect on MRTF-A localization when used at lower concentrations (Figure 1C).

### The application of local force induces actin polymerization and accumulation of MRTF-A in the nucleus

We performed the first set of experiments to measure the impact of mechanical cues on the localization of MRTF-A using magnetic tweezers. A constant force step of 1 nN was applied for 125 s and then released for 125 s, and the cycle repeated six times over a total of 25 min. The localization of MRTF-A-GFP was assessed at the end of each cycle (SI movie 1 and Figure 2.). We tested three populations of cells: cells expressing MRTF-A-GFP only, those co-expressing MRTF-A-GFP and mCherry-actin, and those co-expressing MRTF-A-GFP and LifeAct-mCherry. All three populations exhibited a similar repartition of MRTF-A-GFP before the application of force (SI Figure 4). The final localization of MRTF-A-GFP for the three cell populations is displayed in Figure 2B.

**Figure 4.**
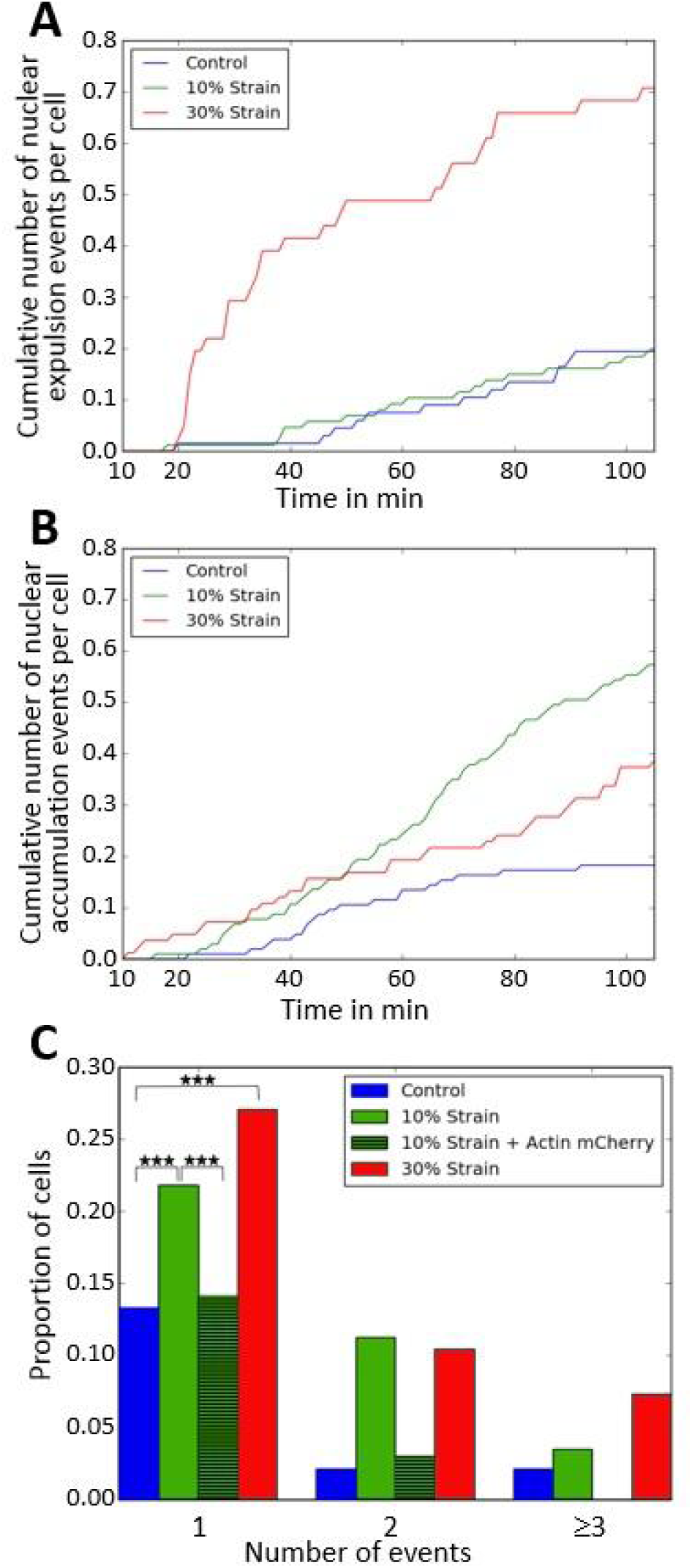
Cumulative number of nuclear accumulation and expulsion events. A.Cumulative rate of nuclear expulsion events, counted as the number of events per observed cells. A nuclear expulsion event corresponds to the observation of a cell changing from the C to the H or N state or from the H to N state. B.Cumulative rate of nuclear accumulation events, counted as the number of events per observed cells. A nuclear accumulation event corresponds to the observation of a cell changing from the N to the H or C state or changing from the H to C state. C.The proportion of cells experiencing 1, 2, or 3 or more changes in main MRTF-A-GFP localization during the 110 min of observations under control (no strain) (143 cells), 10% strain (145 cells expressing MRTF-A-GFP only, 176 cells co-expressing MRTF-A-GFP and mCherry-actin), or 30% strain (101 cells).

In the absence of actin or actin-binding protein overexpression, mechanically stimulated cells (“+*F*” in Fig. 2) showed a marked increase in the nuclear localization of MRTF-A-GFP, whereas the localization of MRTF-A-GFP did not significantly change in control non-stimulated cells. Changing the F/G-actin equilibrium interfered with the nuclear re-localization of MRTF-A-GFP under mechanical stress. Overexpression of actin completely abolished the mechanically-induced nuclear accumulation of MRTF-A-GFP, with localization similar to that of the un-stimulated control, whereas cells expressing LifeAct-mCherry, known to stabilize F-actin [^37^,^38^], exhibited enhanced nuclear re- localization of MRTF-A-GFP under mechanical stimulation.

This response to force may have been associated with a local increase in the amount of F-actin around the mechanically-stimulated beads (SI movie 2): in cells expressing LifeAct-mCherry, a marker of F-actin, the mean fluorescence intensity per pixel in the red channel was measured in a 5-µm-wide ring-shaped area at the periphery of each bead (see SI Fig. 4) and normalized to the mean intensity per pixel for the entire cell. This ratio increased by 80% on average throughout the complete force application process (Figures 2C and D). Indeed, experiments with optical tweezers on the same cell type have already shown a similar effect [^39^].

Cells expressing LifeAct-mCherry displayed increased nuclear localization of MRTF-A-GFP, even without mechanical stimulation (“*F*=0” in Figure 2B). This may be due to the stabilizing effect of LifeAct on actin filaments [^37^,^38^]. The presence of LifeAct thus favors the nuclear localization of MRTF-A-GFP, similarly to SiR-actin, another stabilizer of actin filaments, as described in the previous paragraph.

Overall, these experiments showed that the local application of a force on myoblasts triggered nuclear accumulation of MRTF-A-GFP, which correlated with the local polymerization of actin. Stabilizing actin filaments enhanced this effect, whereas over-expressing actin abolished it. These observations are consistent with this accumulation being due to a shortage of G-actin in the cytoplasm following actin polymerization induced by mechanical stress.

### The stationary localization of MRTF-A in a population of unstimulated cells is maintained through a dynamic equilibrium

In a second series of experiments, the cells were seeded onto stretchable fibronectin-coated PDMS disks (see Materials and Methods) and cultured for 24 h under standard culture conditions (10% serum). The localization of MRTF-A-GFP in a representative population of cells was then tracked over time: for each experiment, images of areas containing at least one cell expressing MRTF-A-GFP were recorded, with a new image of each area taken every 5 to 10 min, allowing each cell to be tracked over time. The fraction of cells in the three states of major MRTF-A-GFP localization (C, H, or N) were thus followed over time (Figure 3). Although cells were tracked over time, not all cells were observed at the same time. The curves result from several independent experiments, whereas several fields of view were tracked over time during a given experiment. Each point thus gives the state of the cells observed at time *t* +/-3 min after stretching, and the curves start at *t* = 3 min.

The changes in the state of major MRTF-A-GFP localization (C, H or N) in live experiments was tracked for individual cells to gain insight into the dynamics of the cell population. The events were divided into two categories: MRTF-A-GFP nuclear accumulation events (changes in the state of a cell from “C” to “H” or “N’ and from “H” to “N”) and MRTF-A-GFP nuclear expulsion events (changes in the state of a cell from “N” to “H” or “C’ and from “H” to “C”). When normalized, the number of events is divided by the number of cells that can undergo the event, *i.e.* “C” and “H” cells for accumulation events and “H” and “N” cells for expulsion events. The cumulative numbers of events per cell are shown in Figure 4.

The mean localization of MRTF-A-GFP within the cell populations remained roughly stable over time for unstretched cells, with 55% ± 5% of the cells having MRTF-A-GFP mainly in the cytoplasm (Figure 3). This stability resulted from a dynamic equilibrium, as shown in Figure 4: some cells accumulated MRTF-A-GFP in the nucleus over time, whereas MRTF-A-GFP exited the nucleus for others. The rate of nuclear expulsion events was very low and constant over time, one event per approximately 400 cells/min. The rate of nuclear accumulation events was slightly more variable, but with the same mean value, and the population state stayed stable over time.

### A moderate strain induces both short- and long-term nuclear accumulation of MRTF-A-GFP

In stretching experiments, PDMS disks seeded with cells were subjected to static strain applied at time *t* = 0 and held constant over time for the duration of the experiment, typically 110 min. Two different levels of strain were tested, a moderate strain, consisting of a 10% increase in the area of the stretchable substrate, and a higher, 30%, strain. The proportions of cells with mainly “C, “N”, or “H” MRTF-A-GFP when subjected to a constant strain of either 10 or 30% are shown as a function of time in Figure 3, along with a control experiment for unstrained cells.

For the 10% stretch, there were three phases: (i) a first wave of rapid nuclear accumulation of MRTF-A-GFP (< 6 min after stretching), followed by (ii) partial recovery with a new increase in the number of “C” cells and decrease in the number of “H” and “N” cells, and then (iii) a second slow wave of nuclear accumulation of MRTF-A-GFP(*t* > 12 min). The percentage of cells with predominantly cytoplasmic localization of MRTF-A-GFP decreased significantly from 55% before stretching to 35% after 100 min of stretching.

The rate of nuclear expulsion events was the same as in the control (Figure 4), low and constant over time. On the contrary, the rate of nuclear accumulation events was more than two times higher than in the control between 20 and 60 min after stretching, and increased even more, to one event per approximately100 cells/min after 60 min, before slowing down to one event per approximately 250 cells/min for t > 80 min. There was hence a net balance towards nuclear accumulation at all times (after more than 10 min) (Figure 3A). It is noteworthy that the first rapid nuclear accumulation of MRTF-A-GFP (*t* < 6 min), followed by rapid expulsion (*t* < 10min), occurred over the entire cell population as shown in Figure 3A, but did not appear in the cumulative events shown in Figure 4, as it occurred below the temporal resolution of the experiment (approximately 10 min, which was the interval between two successive observations of the state of the same cell).

### A higher, 30%, strain induces short- and long-term nuclear accumulation, but nuclear expulsion of MRTF-A-GFP at intermediate times

Under conditions of 30% stretching, the percentages of cells in the C and H states at t *t* = 3 min were both approximately 40%, demonstrating rapid nuclear accumulation of MRTF-A-GFP. There were then four time-dependent phases: (i) a rapid increase in the number of “C” cells accompanied by a rapid decrease in the number of “N” cells (*t* < 10 min after stretching), followed by (ii) a rapid decrease in the number of “C” cells accompanied by a rapid increase in the number of “H” cells (10 < t < 15 min), followed in the longer term by (iii) a second slow increase in the number of “C” cells, up to 65%, and finally (iv), a second decrease in the number of “C” cells accompanied by an increase in the number of “H” cells for *t* > 80 min.

Nuclear accumulation of MRTF-A-GFP occurred earlier than in control and 10% stretching experiments, as early as could be observed (10 min after stretching, Figure 4B), and the rate remained almost constant over the entire experiment, with approximately one nuclear accumulation event for 250 cells/min, a rate more than two times higher than in control experiments. This led to the observed decrease in the number of “C” cells between 10 and 15 min (Figure 3B). However, at *t* ≈ 20 min, the rate of nuclear expulsion events rapidly increased to one nuclear expulsion event for 44 cells/min, a rate almost 10 times higher than in control experiments (Figure 4B). This rate remained very high for 20 min and then decreased over the next hour, but remained higher than that of accumulation events (1 for 250 cells/min) until it reached the value of the control experiments for *t* > 80min (1 for 400 cells/min). Hence, the net balance was nuclear accumulation from10 to 20 min, then nuclear expulsion from 20 to 80 min, and finally slow nuclear accumulation at longer time-points (*t* > 80 min).

In conclusion, under a maintained stretch, myoblasts rapidly accumulated MRTF-A-GFP in the nucleus, within a few minutes. This was followed by expulsion of MRTF-A-GFP back into the cytoplasm. The duration and intensity of the expulsion stage depended on the degree of stretch. Under 10% strain, MRTF-A-GFP re-accumulated again in the nucleus from 10 min up to 2 h, whereas under 30% strain, expulsion was stronger and continued for up to 80 min before partial recovery of the nuclear localization of MRTF-A-GFP.

### Both degrees of strain cause a significant increase of re-localization events in the population, which are suppressed by over-expressing actin

There was a clear difference between the total number of change of state events for the control and stretching experiments during the 100 min of an entire experiment (Figure 4C). In control experiments (without strain), a little less than 20% of the cells experienced one or several changes in the main localization of MRTF-A-GFP during the 100 min. The number of events per cell doubled when a 10% strain was applied for 100 min, with approximately 38% of the cells showing a change in the main localization of MRTF-A-GFP at least once during the experiment. Under a 30% applied strain, this proportion increased to 46%. The difference was particularly clear concerning the percentage of cells exhibiting two or more changes in the main localization of MRTF-A-GFP: 6% for the control, 20% for the 10% stretch, and 20% for the 30% stretch.

Overexpression of actin through mCherry-actin completely inhibited nuclear translocation of MRTF-A under mechanical stimulation, as previously observed for the magnetic tweezers experiments: the proportion of cells displaying cytoplasmic localization of MRTF-A-GFP remained constant over time, as in the control experiments, and the number of events was also very similar (Figure 4C), with MRTF-A retained in the cytoplasm with the increasing pool of actin.

### A moderate strain induces actin polymerization, whereas a high strain induces actin depolymerization at intermediate times

We investigated the actin dynamics and correlation between actin polymerization and MRTF-A nuclear accumulation under strain by staining the cells with the live F-actin fluoregenic probe SiR-actin (40) at a concentration of 50nM, a concentration that did not alter the sub-cellular repartition of MRTF-A (cf. Figure 1C). SiR-actin was added to the cell medium and stained all the cells. Its fluorescence was much higher when attached to actin filaments than when in solution (40) and it was used without rinsing, allowing the staining of the newly formed actin filaments.

Live SiR-actin monitoring of stretched cells (Figure 5A) revealed three phases of actin polymerization in response to a 10% strain, bursts at 5 to 18 min and a second at 25 to 35 min, followed by a continuous and marked increase in the level of F-actin beginning approximately 45 min after the start of stretching. The second burst and long-term actin polymerization correlated well with the nuclear accumulation of MRTF-A-GFP (Figure. 5B). The very short-term behavior was less clear, because the time resolution of these experiments was approximately10 min.

**Figure 5.**
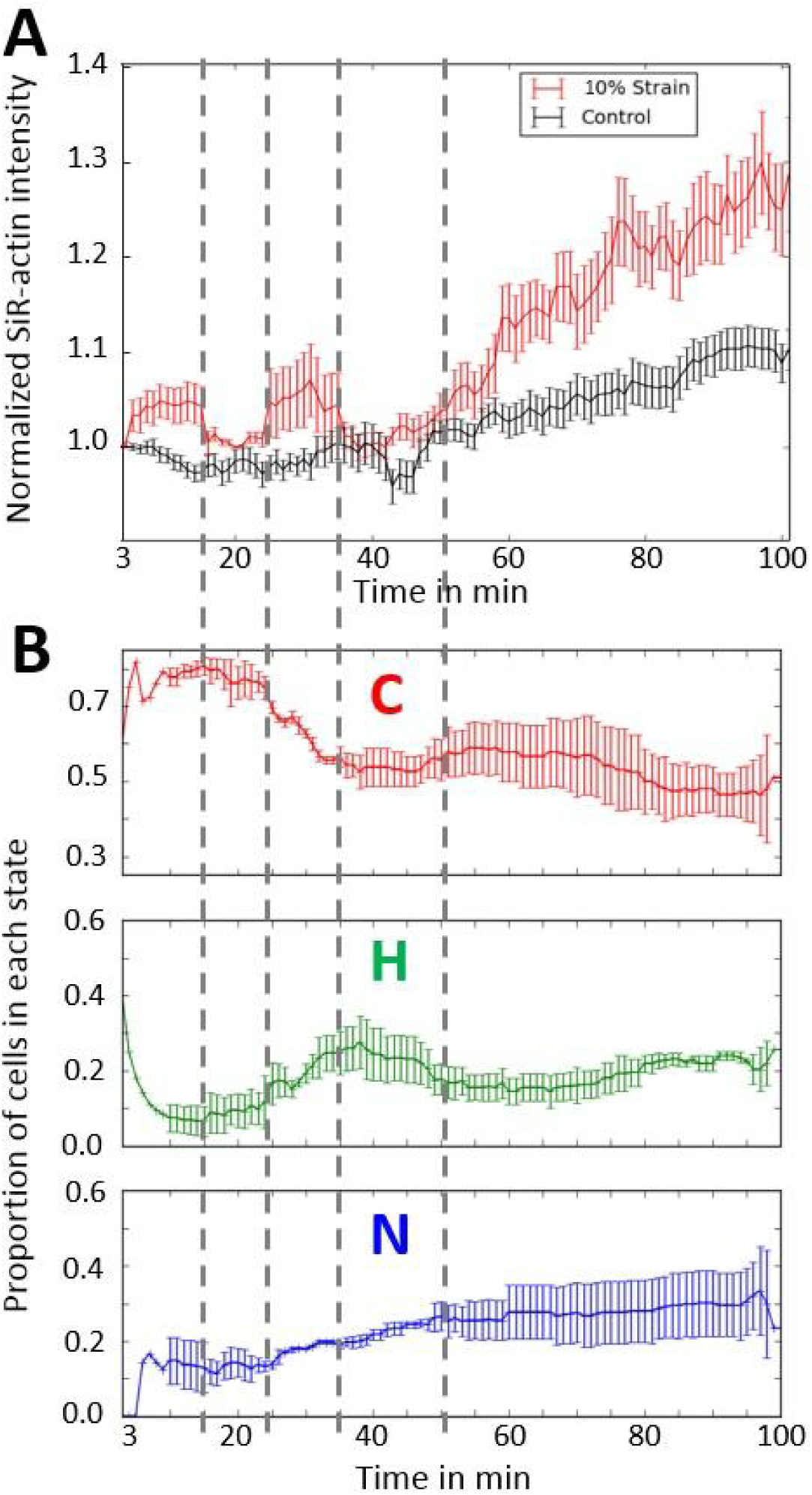
A. Mean value of the SiR-actin intensity relative to the initial fluorescence level in cells submitted to a 10% stretch at t = 0 and under control conditions. 10% strain: n = 3 independent experiments, 93 cells; control: n = 2 independent experiments, 124 cells. B. Evolution of over time of the repartition among the three categories (C, N, and H) of MRTF-A-GFP localization for the same cells as in A.

In the control experiments with SiR-actin, unstretched cells showed a slight decrease in short-term SiR-actin intensity, presumably due to phototoxicity, as well as a small long-term increase, revealing that SiR-actin, a derivative of jasplakinolide, still acted as a stabilizer of actin filaments, even at the very low concentration used here. Figure 5B differs from Figure 3A because the samples differed in two ways: first the use of SiR-actin and second the transfection agent used (Nanofectin for the samples of Figure 3 and Lipofectamine for the samples of Figure 5, see Materials and Methods).

Cells were also stretched at a higher strain, a 30% increase in area, following the same experimental protocol. The cytoskeleton appeared to be incapable of sustaining such a high degree of deformation: SiR-actin intensity started decreasing as soon as 10 min after the start of stretching and continued to decrease for up to 35 min (Figure 6), suggesting damage to the cytoskeleton (also apparent in Figure 7A). Actin depolymerization correlated with the nuclear expulsion of MRTF-A-GFP (Figure 3B). F-actin re-polymerized at longer timepoints, *t* > 35min (see Fig. 6).

**Figure 6.**
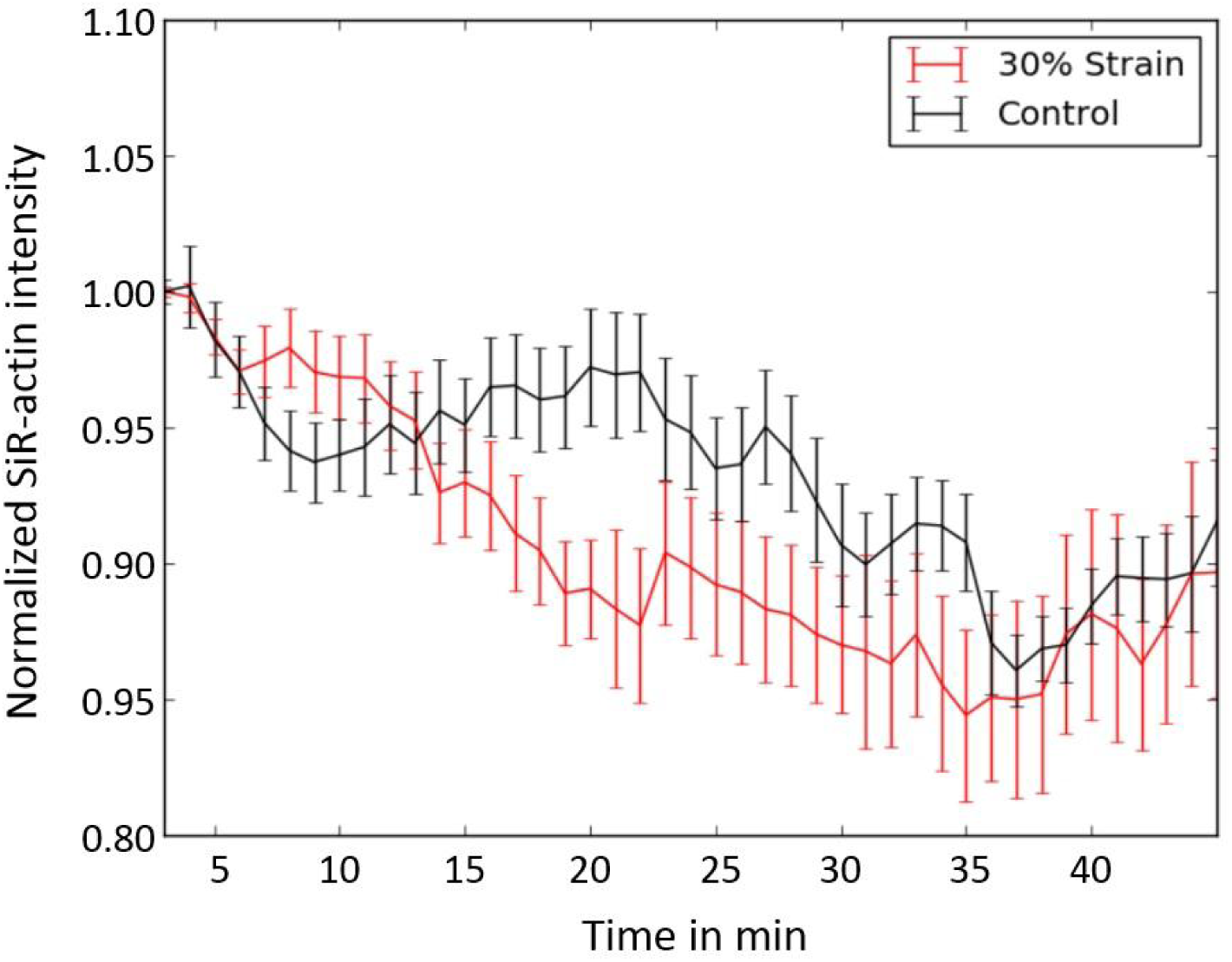
Median SiR-actin intensity relative to the initial fluorescence level. (30% strain: *n* = 4 independent experiments; 108 cells. Control *n* = 2, 53 cells).

**Figure 7.**
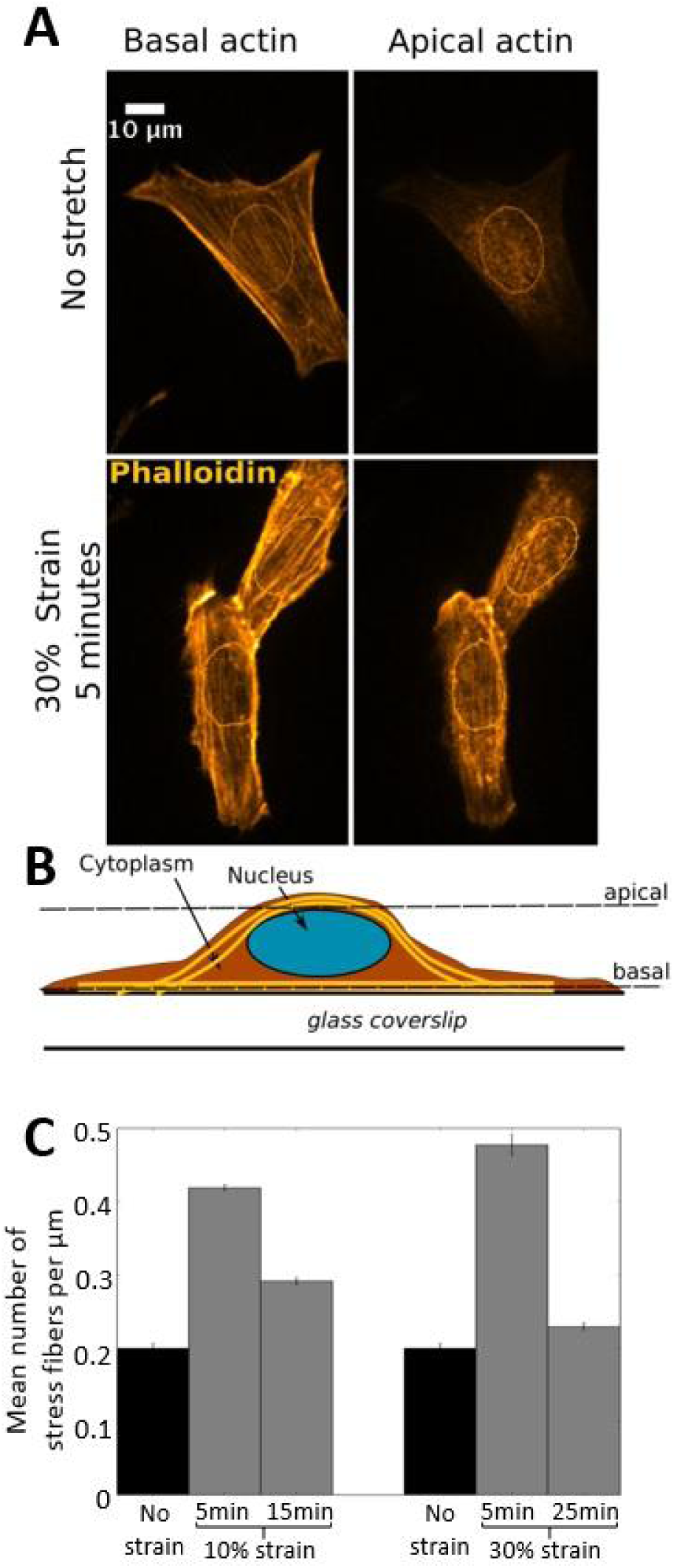
A. Confocal microscopy images of myoblasts in which F-actin was labelled with SiR-actin. top: un-stretched control cells; top left: basal layer showing aligned actin stress fibers; top right: apical actin with un-organized actin structure. bottom: cells after 5 min of 30% strain; bottom left: basal layer with aligned actin stress fibers, bottom right: organized actin cap with aligned stress fibers above the nucleus. The contour of the nuclei was measured from the DAPI signal and drawn on the images. B.Schematic view of a cell, showing the apical and basal levels that were imaged. C.Mean number of stress fibers above the nucleus rescaled by the size of the nucleus (width in µm in the direction perpendicular to the stress fibers). Control: 38 cells; 10% strain - t = 5 min: 66 cells, 10% strain – t = 15 min: 55 cells, 30% strain – t = 5min: 50 cells, 30% strain – t = 25 min: 41 cells.

Overall, these results show that stretching the cells triggered either time-dependent assembly or disassembly of actin filaments, depending on the strain level, and that F-actin assembly correlated with nuclear accumulation of MRTF-A-GFP, whereas F-actin disassembly correlated with cytoplasmic accumulation of MRTF-A-GFP.

### Stress fibers form a transient actin cap above the nucleus minutes after stretching

We fixed samples after 5, 15, or 25 min of stretch to gain insight into the phenomena that occur shortly after the initiation of stretch. SiR-actin was used after fixation to stain F-actin and the cells were observed by confocal microscopy. In the initial state (no stretching), apical F-actin showed no particular structure in most of the cells (Figure. 7A). However, some cells showed parallel and organized stress fibers above the nucleus, a structure known as an actin cap [^40^,^41^]: 45% of the cells had no actin cap and 25% had only one fiber. After 5 min of stretch, nearly all the cells displayed an organized actin cap and the mean number of stress fibers above the nucleus, rescaled by the width of their nuclei in the direction perpendicular to the stress fibers, doubled after 5 min of 10% stretch and more than doubled after 5 min of 30% stretch (Figure 7C). However, such actin reorganization was highly transient: the actin cap returned to its control configuration after 25 min of maintained stretch. Strikingly, this rapid actin polymerization/depolymerization process correlated with the observed short-term nuclear accumulation/expulsion of MRTF-A-GFP (Figure 3A and B). The overall levels of ventral actin stress fibers did not measurably vary after 5 min of 10% stretch.

On the contrary, when cells were stretched by 30%, the ventral stress fibers were damaged (Figure 7B, left panel) and the overall level of F-actin decreased (Figure. 6). This rapid actin polymerization / depolymerization processes again correlated with the observed short-term MRTF-A-GFP nuclear accumulation followed by rapid nuclear expulsion which lasted until the cells recovered and actin polymerization started over, after approximately 35 min of stretch.

Overall, these results show the rapid formation of an actin cap in response to stretch, associated with MRTF-A accumulation in the nucleus, suggesting that the actin cap is involved in the rapid cellular response to strain [^41^].

## Discussion

The actin/SRF/MRTF-A pathway is amongst the mechanosensitive signaling pathways, transducing mechanical signals to gene expression, and is at the center of mechanotransduction in muscles. The mechanistic link between the F/G actin balance and sub-cellular localization of MRTF-A is well documented [^21^-^26^]. The depletion of the G-actin pool by F-actin polymerization has been shown to enable the transport of MRTF-A into the nucleus, resulting in its nuclear accumulation. However, the dynamics of the system are poorly understood.

Here, we conducted studies on C2C12 myoblasts and first confirmed that the sub-cellular distribution of MRTF-A closely correlates with the actin F/G ratio, with excess G-actin favoring cytoplasmic localization and F-actin stabilization favoring nuclear localization. Moreover, we show that a population of myoblasts under standard culture conditions (*i.e.* in medium containing serum) display a dynamic stationary state for the localization of MRTF-A, in which some cells display nuclear accumulation of MRTF-A, whereas others display nuclear expulsion. We then investigated the dynamics of MRTF-A re-localization in response to various kinds and levels of mechanical stimulation, as well as its links with actin polymerization. We show that the application of local mechanical stress through microbeads induces local polymerization of actin within minutes, as previously observed [^39^], and a significant increase in nuclear localization of MRTF-A-GFP after 30 min of the application of force. We also show, for the first time, the rapid but transient nuclear accumulation of MRTF-A in cells subjected to a global stretch, within a few minutes, which correlates with the formation of a peri-nuclear actin cap. This is similar to the recent observations of the rapid and reversible assembly of the nuclear actin network in serum-stimulated fibroblasts [^27^]. However, the F-actin assembly that we observed was peri-nuclear rather than nuclear. Under sustained stretch, we also observed long-term polymerization of ventral actin, which correlated with nuclear accumulation of MRTF-A, lasting for the two hours of our experiments. The time between the observed actin polymerization and MRTF-A nuclear accumulation was below the temporal resolution of our experiments, which was a few minutes, as previously observed [^39^].

In conclusion, we have shown that various types of mechanical stimulation on myoblasts induce nuclear accumulation of MRTF-A, which correlates with actin polymerization in various elements of the cytoskeleton. These phenomena occurred over a wide range of time-scales, from the temporal resolution of the experiments (a few minutes) to their duration (approximately two hours). All our results are consistent with known mechanisms of MRTF-A regulation by G-actin, with a strong correlation between F-actin assembly and the nuclear accumulation of MRTF-A. Furthermore, we demonstrated that nuclear accumulation of MRTF-A after mechanical stimulation is maintained over the long-term, which is probably essential for mechano-transduction.

Future studies will need to investigate the same questions in more physiological systems, such as primary myoblasts and myotubes.

## Author contributions

L.M. designed and carried out experiments, analyzed results and wrote the manuscript. A.S. and S.H. designed experiments, analyzed results, wrote the manuscript and provided financial support.

## Data availability

The datasets generated during and/or analyzed during the current study are available from the corresponding author on request.

## Supplementary information

The strain rate can be easily computed using a simple geometric model. The increase in area depends on the free PDMS sheet radius *R*_*l*_, the post radius *R*_*p*_, and the depth it pushes the sheet down *h*. (see SI Figure. 2). The initial area is *πR*_*l*_^*2*^ and the final area is the sum of the post area *πR*_*p*_^*2*^ and the truncated cone area *π(R*_*p*_*+R*_*l*_*)[(R*_*l*_*-R*_*p*_*)²+h²]*^*1/2*^. Finally, the relative increase in area is:

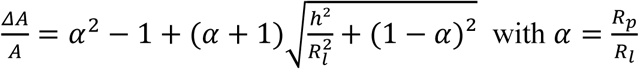

**Figure S1.**
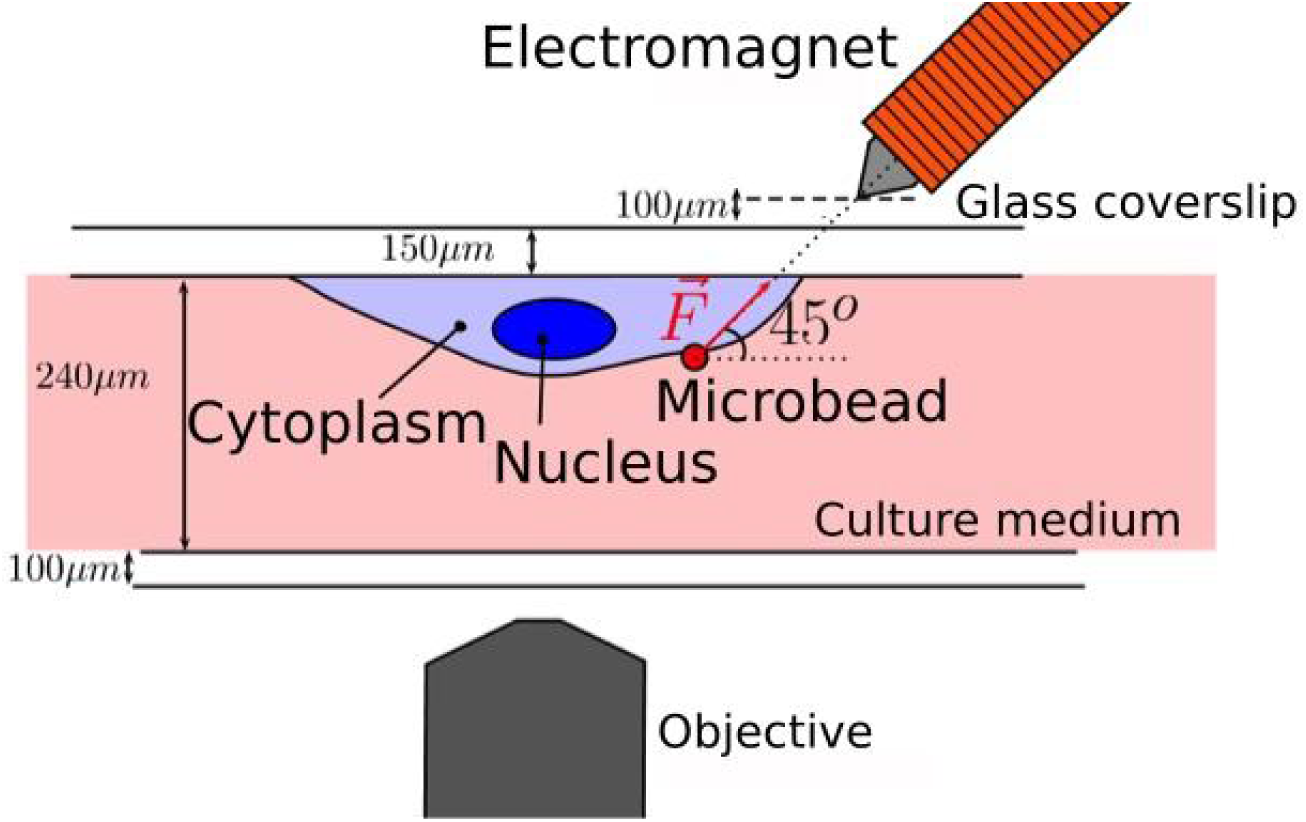
Schematic view of the magnetic tweezers experiment. The cells are upside down on the top coverslip to obtain the shortest possible distance between the tip of the electromagnet and the attached bead. The applied force is estimated to be approximately 1 nN. It is kept constant for 125 s every 250 s for six cycles.

**Figure S2.**
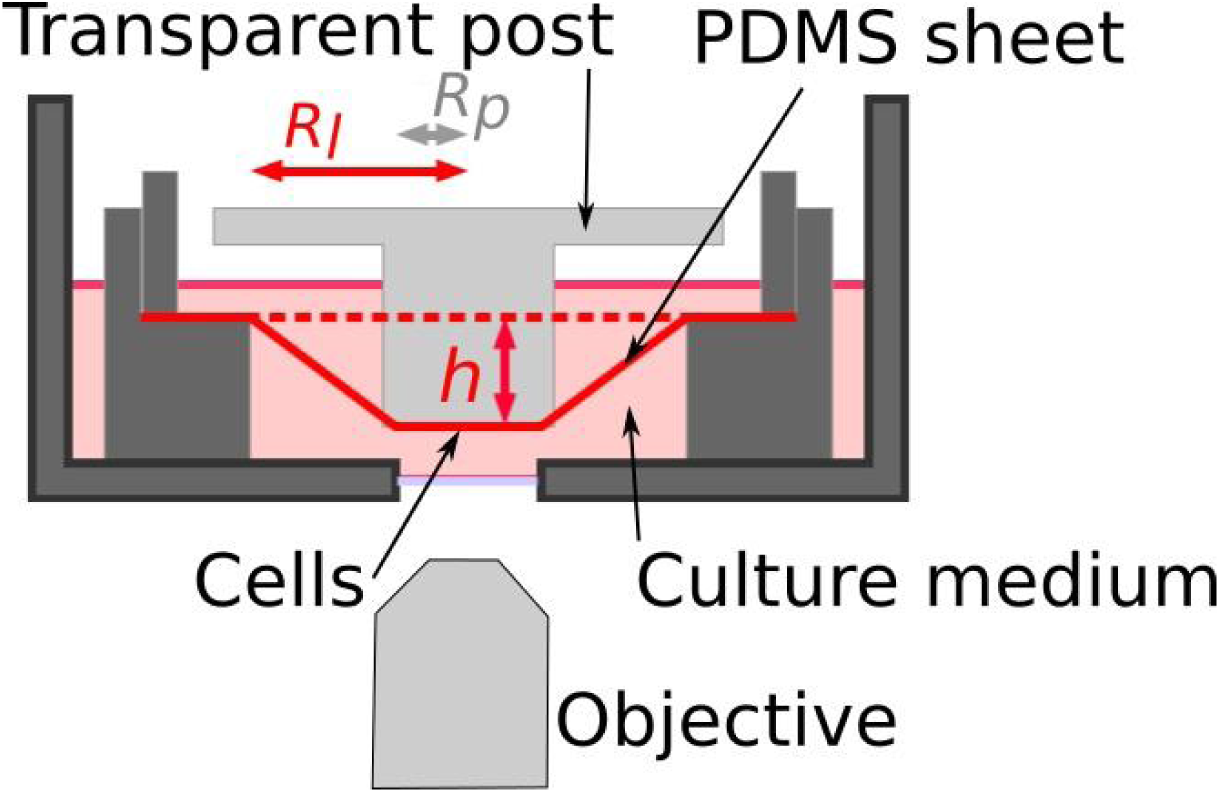
Schematic view of the stretching device. The cells are on the underside of the fibronectin- coated stretched PDMS sheet and are observed from below using an inverted microscope. At time t = 0, the transparent post is pushed down to a depth *h*, which causes the strain.

**Figure S3.**
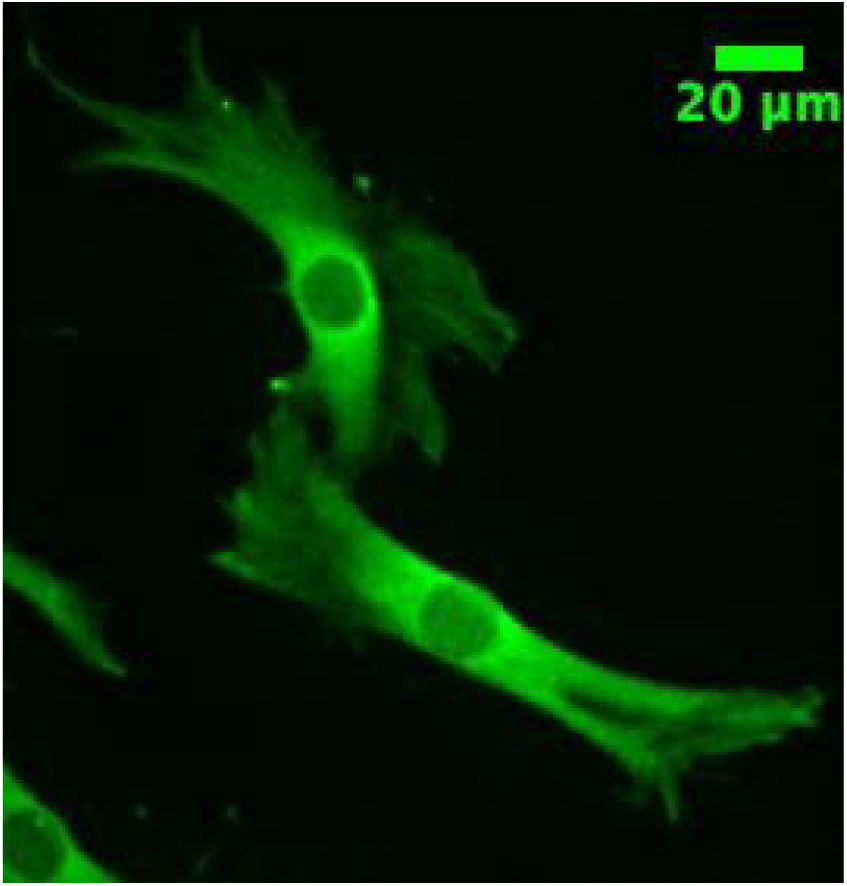
Immunofluorescence images of C2C12 stained with anti-MRTF-A. MRTF-A is mainly cytoplasmic in almost all cells.

**Figure S4.**
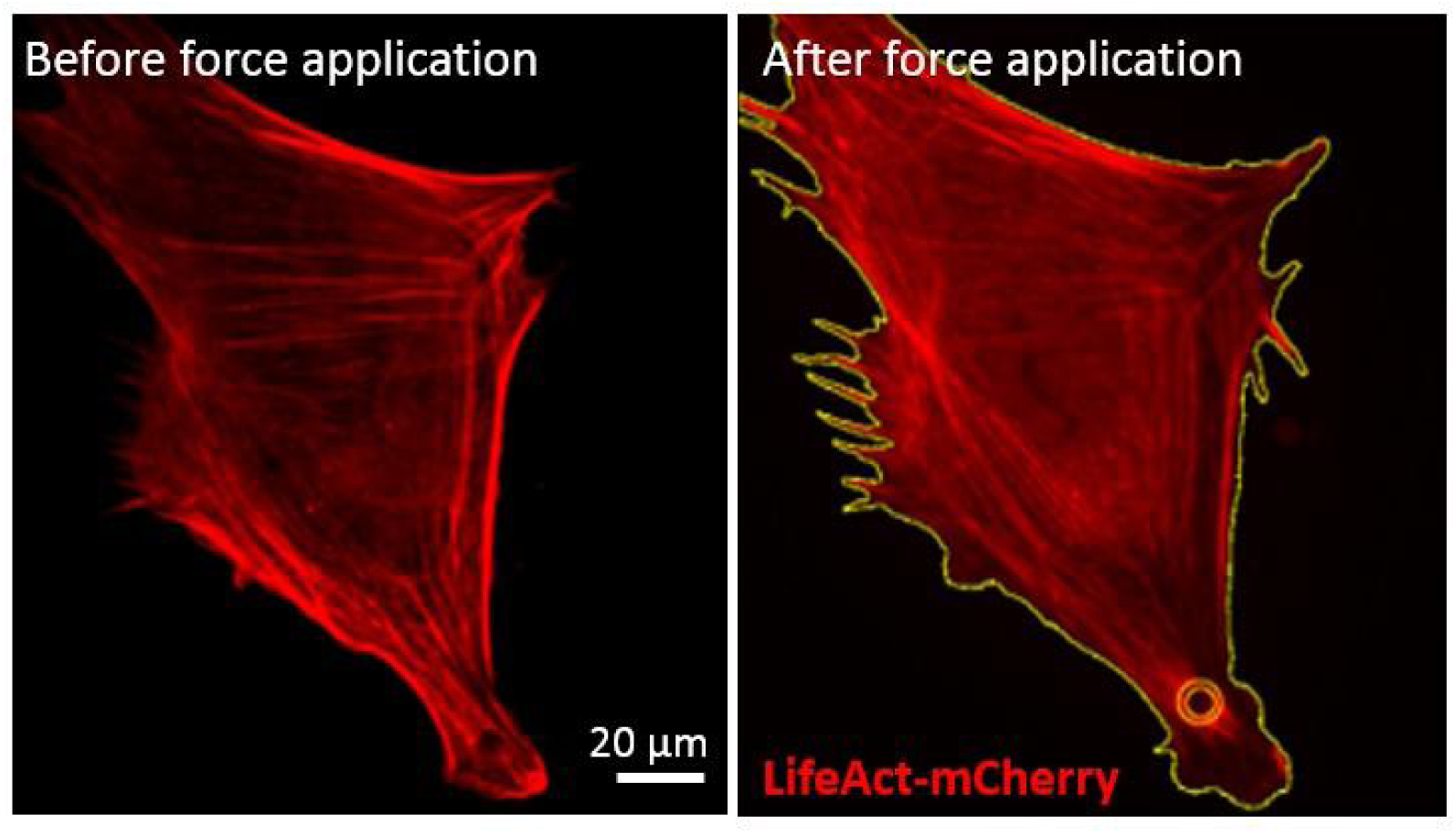
Example of the areas used to assess actin enrichment around the microbead. The mean intensity per pixel in the LifeAct-mCherry channel in a 5-μm wide ring around the bead is compared to that of the whole cell.

## References

1. Janmey, P. A., Wells, R. G., Assoian, R. K. & McCulloch, C. A. From tissue mechanics to transcription factors. Differentiation 86, 112–120 (2013).

2. Iba, T. & Sumpio, B. E. Morphological response of human endothelial cells subjected to cyclic strain in vitro. Microvascular Research 42, 245–254 (1991).

3. Li, S. et al. Requirement for serum response factor for skeletal muscle growth and maturation revealed by tissue-specific gene deletion in mice. Proceedings of the National Academy of Sciences 102, 1082–1087 (2005).

4. Janoštiak, R., Pataki, A. C., Brábek, J. & Rösel, D. Mechanosensors in integrin signaling: The emerging role of p130Cas. European Journal of Cell Biology 93, 445–454 (2014).

5. Blanchoin, L., Boujemaa-Paterski, R., Sykes, C. & Plastino, J. Actin Dynamics, Architecture, and Mechanics in Cell Motility. Physiological Reviews 94, 235–263 (2014).

6. Morita, T., Mayanagi, T. & Sobue, K. Reorganization of the actin cytoskeleton via transcriptional regulation of cytoskeletal/focal adhesion genes by myocardin-related transcription factors (MRTFs/MAL/MKLs). Experimental Cell Research 313, 3432–3445 (2007).

7. Ahmed, W. W. et al. Myoblast morphology and organization on biochemically micro- patterned hydrogel coatings under cyclic mechanical strain. Biomaterials 31, 250–258 (2010).

8. Mendez, M. G. & Janmey, P. A. Transcription factor regulation by mechanical stress. The International Journal of Biochemistry & Cell Biology 44, 728–732 (2012).

9. Sotiropoulos, A., Gineitis, D., Copeland, J. & Treisman, R. Signal-Regulated Activation of Serum Response Factor Is Mediated by Changes in Actin Dynamics. Cell 98, 159–169 (1999).

10. Wang, D.-Z. et al. Potentiation of serum response factor activity by a family of myocardin-related transcription factors. Proceedings of the National Academy of Sciences 99, 14855–14860 (2002).

11. Cen, B. et al. Megakaryoblastic Leukemia 1, a Potent Transcriptional Coactivator for Serum Response Factor (SRF), Is Required for Serum Induction of SRF Target Genes. Molecular and Cellular Biology 23, 6597–6608 (2003).

12. Kalita, K., Kharebava, G., Zheng, J.-J. & Hetman, M. Role of Megakaryoblastic Acute Leukemia-1 in ERK1/2-Dependent Stimulation of Serum Response Factor-Driven Transcription by BDNF or Increased Synaptic Activity. Journal of Neuroscience 26, 10020–10032 (2006).

13. Kalita, K., Kuzniewska, B. & Kaczmarek, L. MKLs: Co-factors of serum response factor (SRF) in neuronal responses. The International Journal of Biochemistry & Cell Biology 44, 1444–1447 (2012).

14. Gerber, A. et al. Blood-Borne Circadian Signal Stimulates Daily Oscillations in Actin Dynamics and SRF Activity. Cell 152, 492–503 (2013).

15. Gomez, E. W., Chen, Q. K., Gjorevski, N. & Nelson, C. M. Tissue geometry patterns epithelial–mesenchymal transition via intercellular mechanotransduction. Journal of Cellular Biochemistry 110, 44–51 (2010).

16. Connelly, J. T. et al. Actin and serum response factor transduce physical cues from the microenvironment to regulate epidermal stem cell fate decisions. Nature Cell Biology 12, 711–718 (2010).

17. Selvaraj, A. & Prywes, R. Megakaryoblastic Leukemia-1/2, a Transcriptional Co-activator of Serum Response Factor, Is Required for Skeletal Myogenic Differentiation. J. Biol. Chem. 278, 41977–41987 (2003).

18. Esnault, C. et al. Rho-actin signaling to the MRTF coactivators dominates the immediate transcriptional response to serum in fibroblasts. Genes & Development 28, 943–958 (2014).

19. Guerci, A. et al. Srf-Dependent Paracrine Signals Produced by Myofibers Control Satellite Cell-Mediated Skeletal Muscle Hypertrophy. Cell Metabolism 15, 25–37 (2012).

20. Collard, L. et al. Nuclear actin and myocardin-related transcription factors control disuse muscle atrophy through regulation of Srf activity. Journal of Cell Science 127, 5157–5163 (2014).

21. Miralles, F., Posern, G., Zaromytidou, A.-I. & Treisman, R. Actin dynamics control SRF activity by regulation of its coactivator MAL. Cell 329–342 (2003).

22. Vartiainen, M. K., Guettler, S., Larijani, B. & Treisman, R. Nuclear Actin Regulates Dynamic Subcellular Localization and Activity of the SRF Cofactor MAL. Science 316, 1749–1752 (2007).

23. Mouilleron, S., Guettler, S., Langer, C. A., Treisman, R. & McDonald, N. Q. Molecular basis for G-actin binding to RPEL motifs from the serum response factor coactivator MAL. The EMBO Journal 27, 3198–3208 (2008).

24. Guettler, S., Vartiainen, M. K., Miralles, F., Larijani, B. & Treisman, R. RPEL Motifs Link the Serum Response Factor Cofactor MAL but Not Myocardin to Rho Signaling via Actin Binding. Mol. Cell. Biol. 28, 732–742 (2008).

25. Hirano, H. & Matsuura, Y. Sensing actin dynamics: Structural basis for G-actin-sensitive nuclear import of MAL. Biochemical and Biophysical Research Communications 414, 373–378 (2011).

26. Pawłowski, R., Rajakylä, E. K., Vartiainen, M. K. & Treisman, R. An actin-regulated importin α/β-dependent extended bipartite NLS directs nuclear import of MRTF-A. The EMBO Journal 29, 3448–3458 (2010).

27. Baarlink, C., Wang, H. & Grosse, R. Nuclear Actin Network Assembly by Formins Regulates the SRF Coactivator MAL. Science 340, 864–867 (2013).

28. Lundquist, M. R. et al. Redox Modification of Nuclear Actin by MICAL-2 Regulates SRF Signaling. Cell 156, 563–576 (2014).

29. Scharenberg, M. A. et al. TGF- -induced differentiation into myofibroblasts involves specific regulation of two MKL1 isoforms. Journal of Cell Science 127, 1079–1091 (2014).

30. Zhao, X.-H. et al. Force activates smooth muscle -actin promoter activity through the Rho signaling pathway. Journal of Cell Science 120, 1801–1809 (2007).

31. Albinsson, S., Nordström, I. & Hellstrand, P. Stretch of the Vascular Wall Induces Smooth Muscle Differentiation by Promoting Actin Polymerization. J. Biol. Chem. 279, 34849–34855 (2004).

32. Staus, D. P., Blaker, A. L., Taylor, J. M. & Mack, C. P. Diaphanous 1 and 2 Regulate Smooth Muscle Cell Differentiation by Activating the Myocardin-Related Transcription Factors. Arteriosclerosis, Thrombosis, and Vascular Biology 27, 478–486 (2007).

33. Kuwahara, K. et al. Myocardin-Related Transcription Factor A Is a Common Mediator of Mechanical Stress- and Neurohumoral Stimulation-Induced Cardiac Hypertrophic Signaling Leading to Activation of Brain Natriuretic Peptide Gene Expression. Mol. Cell. Biol. 30, 4134–4148 (2010).

34. Huang, X. et al. Matrix Stiffness–Induced Myofibroblast Differentiation Is Mediated by Intrinsic Mechanotransduction. American Journal of Respiratory Cell and Molecular Biology 47, 340–348 (2012).

35. Chan, M. W. C., Chaudary, F., Lee, W., Copeland, J. W. & McCulloch, C. A. Force-induced Myofibroblast Differentiation through Collagen Receptors Is Dependent on Mammalian Diaphanous (mDia). J. Biol. Chem. 285, 9273–9281 (2010).

36. Iyer, K. V., Pulford, S., Mogilner, A. & Shivashankar, G. V. Mechanical Activation of Cells Induces Chromatin Remodeling Preceding MKL Nuclear Transport. Biophysical Journal 103, 1416–1428 (2012).

37. Courtemanche, N., Pollard, T. D. & Chen, Q. Avoiding artefacts when counting polymerized actin in live cells with LifeAct fused to fluorescent proteins. Nature Cell Biology 18, 676–683 (2016).

38. Honing, H. S. van der, Bezouwen, L. S. van, Emons, A. M. C. & Ketelaar, T. High expression of Lifeact in Arabidopsis thaliana reduces dynamic reorganization of actin filaments but does not affect plant development. Cytoskeleton 68, 578–587 (2011).

39. Icard-Arcizet, D., Cardoso, O., Richert, A. & Hénon, S. Cell Stiffening in Response to External Stress is Correlated to Actin Recruitment. Biophysical Journal 94, 2906–2913 (2008).

40. Khatau, S. B. et al. A perinuclear actin cap regulates nuclear shape. Proceedings of the National Academy of Sciences 106, 19017–19022 (2009).

41. Chambliss, A. B. et al. The LINC-anchored actin cap connects the extracellular milieu to the nucleus for ultrafast mechanotransduction. Scientific Reports 3, (2013).

